# A Framework for Collective Behavior in Plant Inspired Growth-Driven Systems, Based on Kinematics of Allotropism

**DOI:** 10.1101/566364

**Authors:** Renaud Bastien, Amir Porat, Yasmine Meroz

## Abstract

A variety of biological systems are not motile, but sessile in nature, relying on growth as the main driver of their movement. Groups of such growing organisms can form complex structures, such as the functional architecture of growing axons, or the adaptive structure of plant root systems. These processes are not yet understood, however the decentralized growth dynamics bear similarities to the collective behavior observed in groups of motile organisms, such as flocks of birds or schools of fish. Equivalent growth mechanisms make these systems amenable to a theoretical framework inspired by tropic responses of plants, where growth is considered implicitly as the driver of the observed bending towards a stimulus. We introduce two new concepts related to plant tropisms: point tropism, the response of a plant to a nearby point signal source, and *allotropism*, the growth-driven response of plant organs to neighboring plants. We first analytically and numerically investigate the 2D dynamics of single organs responding to point signals fixed in space. Building on this we study pairs of organs interacting *via* allotropism, *i.e.*each organ senses signals emitted at the tip of their neighbor and responds accordingly. In the case of local sensing we find a rich phase space. We describe the different phases, as well as the sharp transitions between them. We also find that the form of the phase space depends on initial conditions. This work sets the stage towards a theoretical framework for the investigation and understanding of systems of interacting growth-driven individuals.

## I. INTRODUCTION

In the past couple of decades, there has been an increasing interest in the investigation of collective behavior in systems composed of motile self-propelled entities. Examples of collective behavior are widespread in biological systems, including colonies of bacteria [1, 2], motile cells [3, 4], swarms of insects [5, 6], schools of fish [7, 8], or flocks of birds [9, 10], as well as physical systems including nematic fluids [11, 12], shaken metallic rods [13–16], and nano-swimmers [17]. Despite the diversity of these systems, generalized minimal models have shown that collective behavior may emerge as a results of local interactions between individuals [7, 18–27]. Indeed these models are able to reproduce complex behavior using similar minimal rules. For example in the archetypal Vicsek model [28] a particle assumes the average direction of the particles in its close neighborhood, while in the Couzin model [24] an individual is attracted to neighboring individuals and aligns to them. These simple systems all exhibit complex collective behavior, as well as transitions between disorganized and organized states. The sensory processes involved in biological systems are taken into account by introducing specific attraction, repulsion and alignment interactions [18, 21, 24]. Furthermore this understanding is at the basis of the field of swarm robotics [29].

However a variety of biological systems are not motile, but sessile in nature, relying on movement resulting from continuous growth in the direction of environmental stimuli. For example, in their search for nutrients plant roots change their morphology by growing differentially. In many cases, groups of such growing organisms form striking structures that provide a biological function or are beneficial for the survival of the group, as in the case of plant root systems that adapt their structure according to water content [30, 31]. The manner in which these complex structures form is not yet understood, however it has been suggested that plant roots exhibit collective behavior [32–34]. This is in line with evidence for interactions between individual plant organs, mediated by complex signaling [35–37].

The few currently-available models of interacting growth-driven systems [32] consider the dynamics of the tip alone, disregarding the fundamental implications of growth, namely the accumulation of past trajectories of individuals, which couples space and time, and constrains possible future trajectories. Furthermore, in elongating organs, changes of curvature at the base have a long-range effect on the position and orientation of the apical tip [38].

The goal of this work is to provide a general framework for the study of collective dynamics in interacting systems where movement is due to growth. We will build on the mathematical description of the kinematics of the growth-driven response of plants towards distant external stimuli, called tropisms, and generalize this to the case of a stimulus emanating from a neighboring plant organ. We will investigate the dynamics of a single organ in response to a static point stimulus, where perception is either local or at the tip alone. Finally, we will consider pairwise interactions, where two organs sense each other and respond accordingly, yielding a complex dynamical phase space. Though we refer to plants throughout the paper, the results are general and can be applied to any similar growth-driven system.

## II. A MINIMAL MODEL FOR ALLOTROPISM: THE GROWTH-DRIVEN REORIENTATION OF PLANTS IN RESPONSE TO DETECTED NEIGHBORING PLANTS

Growth-driven movements in plants can be broadly divided into two main groups, nastic movements and tropisms [39]. Nastic movements encompass all movements that are not oriented towards a stimulus, *e.g.* the circadian opening and closing of leaves. Tropisms are the response of plant organs to a distant directional stimulus, *e.g.* shoots grow towards a source of light (phototropism), and grow away from the direction of gravity (gravitropism).

As mentioned in the Introduction, collective dynamics of plant organs are expected to emerge as a result of local plant-plant dynamical interactions, where for example one plant root senses a signal emanating from a neighboring root and, similarly to common tropic responses, reorients its growth according to the direction of the stimulus. Indeed plants have the ability to perceive the nature and intensity of the interactions with neighboring plants through diverse signals, transmitted either above or below ground [40–42]. These signals can be classified as: (i) indirect signals, corresponding to environmental factors modified by the presence of neighbors, such as light and uptake of nutrients; and (ii) direct signals, corresponding to molecules directly produced by neighboring plants, such as aerial volatile organic compounds (VOCs), soluble root exudates, or microbiome and other intermediates. Furthermore, there are observations of self-recognition in plants [43], as well as recognition of kin [44–46]. However depending on the type of interaction, it is not always clear where the sensing occurs, or where the signal is emitted.

Due to the growth-driven nature of this interaction, we name it *allotropism*, from the greek word for *other* or *another*, *α*́*λλoς* (állos). Accordingly, it is natural to describe the kinematics of this response building on current mathematical models describing regular tropic responses.

Here we adopt a geometrical framework we recently developed for the study of plant tropisms [47, 48]. Though the model is general, making minimal assumptions about the building blocks of the system, it successfully accounts for the experimentally observed gravitropic kinematics of different organs from 11 angiosperms. The model also accounts for the kinematics of growing organs [49] in 3D [50], and has also been generalized to take into account time varying stimuli [51]. However, it is limited to the case of distant stimuli yielding a parallel vector field, such as gravity or sunlight, which approach any point along a plant from the same angle, and at the same intensity. In other words the affecting stimulus is invariant per translation and rotation in the plane perpendicular to the direction of the stimulus.

However when considering interacting neighboring plant organs, such as roots, the signal emanates from a nearby point source, and the direction in which the signal is perceived by one organ is going to change dynamically as the organs move in space. Hence the translational and rotational invariances are no longer always valid throughout the dynamics. Without loss of generality we can assume that the signal emanates from the apical part of the roots, and can be sensed either locally along the root, or apically, *i.e.*at the tip alone. The description of a tropic response to a stimulus emanating from a single point constitutes the basis of the model, and in what follows we will term it *point tropism*. Now the system is only invariant per rotation around the point stimulus. Furthermore, for the sake of simplicity we will consider growth inherently, as the driver of the tropic movement, focusing on the limit where elongation is negligible compared to the timescale of the tropic response (non-elongating organs). This will provide an intuitive understanding of the system, which can be easily transferred in future to elongating organs as shown in [49].

Given the geometrical definitions illustrated in Fig. 1a, and following the *AC* model for the regular tropic response of plants [47, 48], the relation between perception and movement in point tropism can be described by the following equation:

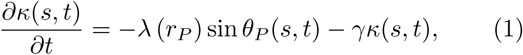

where 0 ≤ *s* ≤ *L* is the curvilinear abscissa along the organ of size *L*, and *θ*(*s, t*) is the angle that the organ forms with the vertical at point *s* and time *t*. The curvature *κ*(*s, t*) is the spatial rate of change of *θ*(*s, t*) along *s*, 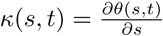. *θ*_*P*_(*s,t*) (*s, t*) is the angle at which the stimulus reaches an element of the organ at point *s* at time *t*, defined as the difference between the angle of the organ and that of the signal point, as illustrated in Fig. 1a:

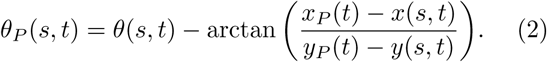

**FIG. 1.**
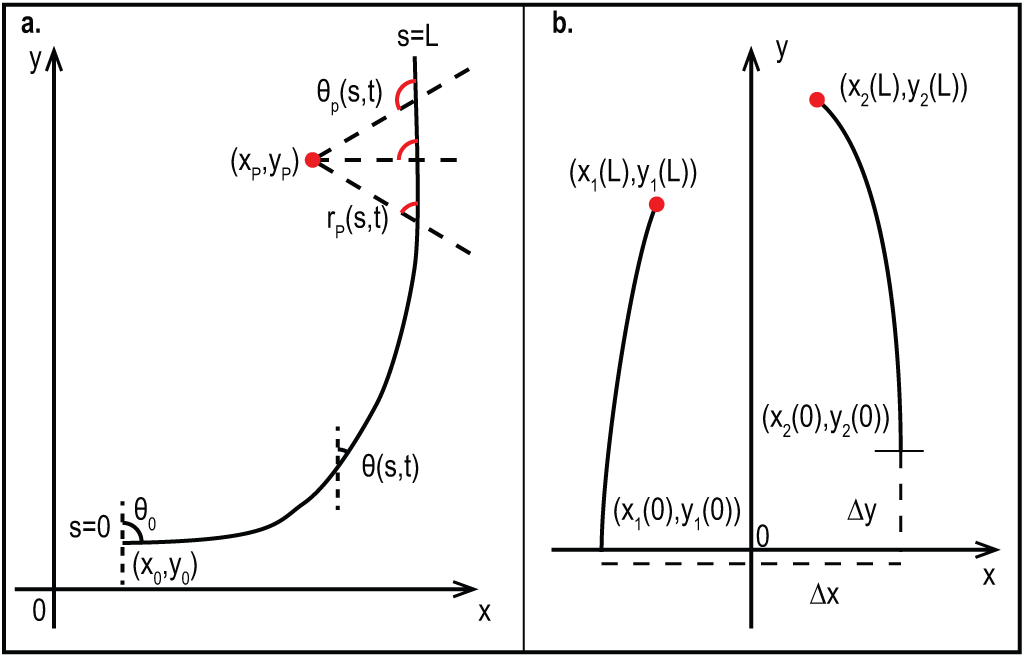
Geometrical definitions. (a) Geometry of point tropism. The median line of an organ is in a plane defined by the coordinates (*x, y*), and described by the parameter *s* which runs along the organ taking the value *s* = 0 at the base, and *s* = *L* at the apical tip, equal to the total length. At each point *s* an angle *θ*(*s, t*) is defined with reference to the direction *y*. The base of the organ is clamped at (*x*(*s* = 0*, t*)*, y*(*s* = 0*, t*)) = (*x*_0_*, y*_0_), at an angle *θ*(*s* = 0*, t*) = *θ*_0_. The position of the point signal is marked (*x*_*P*_, *y*_*P*_). At each point *s* of the organ, the distance to the point signal is *r*_*P*_(*s, t*) while the orientation of this element with respect to the point signal is *θ*_*P*_(*s, t*). (b) Geometry of two interacting organs based on allotropism, the growth-driven response of a plant organ to another. The bases of the left and right organs are clamped at (*x*_1_(0)*, y*_1_(0)) = (− ∆*x/*2, 0) and (*x*_2_(0)*, y*_2_(0)) = (∆*x/*2, ∆*y*) respectively. The positions of the signaling apical tips are then (*x*_1_(*L*)*, y*_1_(*L*)) and (*x*_2_(*L*)*, y*_2_(*L*)). Each organ senses the signaling tip of the neighboring organ.

Here (*x*_*P*_(*t*), *y*_*P*_(*t*)) is the position of the point signal and (*x*(*s, t*)*, y*(*s, t*)) is the spatial position of an element of the organ at point *s*. In contrast to the AC model for regular tropisms such as gravitropism, the perceived angle is not constant along the organ even when the organ is straight, as illustrated in Fig. 1a, and the closer the point stimulus is, the greater the variance along the organ. For comparison, in the AC model Eq. 2 is replaced by *θ*(*s,t*) − *θ*_sig_.

*λ*(*rP*) represents the sensitivity of the plant organ to the signal, and may depend on the distance between the sensing organ element and the signal point

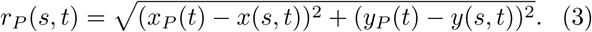

Attractive responses, where the organ bends towards the stimulus, are described by *λ >* 0, and repulsion with *λ <* 0. Here we will consider a constant value, focusing on attractive responses, since repulsive responses are more trivial, as we will discuss later. In line with biological observations [52] the signal is weighed by a sine function and cannot be linearized as has been done previousely for the tropic case [47, 48] since angles cannot be assumed to be small.

Lastly *γ* represents the proprioceptive sensitivity, associated with the ability of plants to perceive their own curvature and to respond accordingly to remain straight. Intuitively, a higher *γ* represents stronger postural control, where the organ resists bending much in response to a stimulus.

The intuition behind Eq. 1 lies in recognizing that the right-hand-side expresses sensing of external signals, represented by the term *λ* sin(*θ*_*P*_(*s, t*)), and internal signals associated with proprioception, *γκ*(*s, t*). On the other hand the left-hand-side represents the growth-driven response represented by the change in curvature 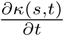.

We note that Eq. 1 assumes that sensing of the signal occurs locally along the whole organ, however in cases where sensing occurs at the tip alone, termed *apical* sensing, the sensing term is replaced with −*λ* sin(*θ*_*P*_(*L, t*)). Furthermore, Eq. 1 holds within a zone of length *L*_gz_ from the tip, *i.e.*0 ≤ *s* ≤ *L* − *L*_gz_, where growth occurs. Outside of the growth zone no response is possible, and Eq. 1 is reduced to 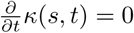. Initial conditions assume that at time *t* = 0 the organ is straight, *κ*(*s, t* = 0) = 0, placed at some angle *θ*_0_ from the vertical *θ*(*s, t* = 0) = *θ*_0_, and clamped at the base, i.e. *θ*(*s* = 0*, t*) = *θ*_0_.

Following dimensional analysis carried out in the AC model [47], we derive characteristic length and time scales. The length scale is identified by considering the steady state, and substituting 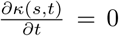 in Eq. 1 while also taking sin *θ*_*P*_ = 1. This yields the maximal curvature value, also termed the *convergence length L*_*c*_ = 1*/κ*_max_ = *γ/λ*. The time scale of the problem is associated with the time it takes for the organ to reach its steady state, termed the *convergence time* and defined as *T*_*c*_ = 1*/γ*. The ratio between the convergence length and the length of the organ introduces a dimensionless number *Z* [53], which describes the balance between the sensitivity to external stimuli and proprioception, and is linearly related to the maximal curvature:

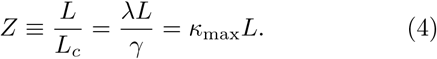

As opposed to the AC model [47], here the angle of the stimulus is not constant, but rather obeys a non-linear dependence on space and time, as expressed by *θ*_*P*_(*s, t*) in Eq. 2. This does not allow analytical calculations, and requires numerical simulations. We note that for distant stimuli this model coincides with that for regular tropisms.

Now that the mathematical model is in place, we will proceed to describe the dynamics of a single organ as a response to a point signal fixed in space, both for apical and local perception.

## III. SINGLE ORGAN DYNAMICS WITH FIXED POINT STIMULUS

We now consider the response of a single organ to a point stimulus fixed in space, focusing on the case of a constant attractive interaction. In what follows we perform out analysis with values of *Z* in the biologically relevant range 1 ≤ *Z* ≤ 10 [47], where the limits *Z* = 1 and *Z* = 10 represent the extreme cases of strong and weak postural control. Numerical simulations are based on a simple forward integration Verlet algorithm, identical to what was used in previous work [47, 48]. A steady state is defined as the configuration at which 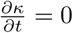 and the organ stops moving.

We first consider the case of apical sensing, where only the apical tip can perceive the direction of the signal, substituting *θ*_*P*_(*L, t*) in Eq. 1. This means that the whole organ responds to the same signal, and thus the dynamics are expected to converge smoothly to the steady state, and the curvature is constant along the organ [48]. Given an initially straight and vertical organ, with *κ*(*s*, 0) = 0 and *θ*(*s*, 0) = 0, the position of the apex is then limited to a set of possible positions, and is calculated analytically in the Supplementary Material (SM). Fig. 2a shows the steady state configurations of organs with different values of *Z*, with point stimuli placed at different positions (represented by a red dot). The configurations are overlaid on the said trajectory of all possible apex positions, plotted in solid black.

**FIG. 2.**
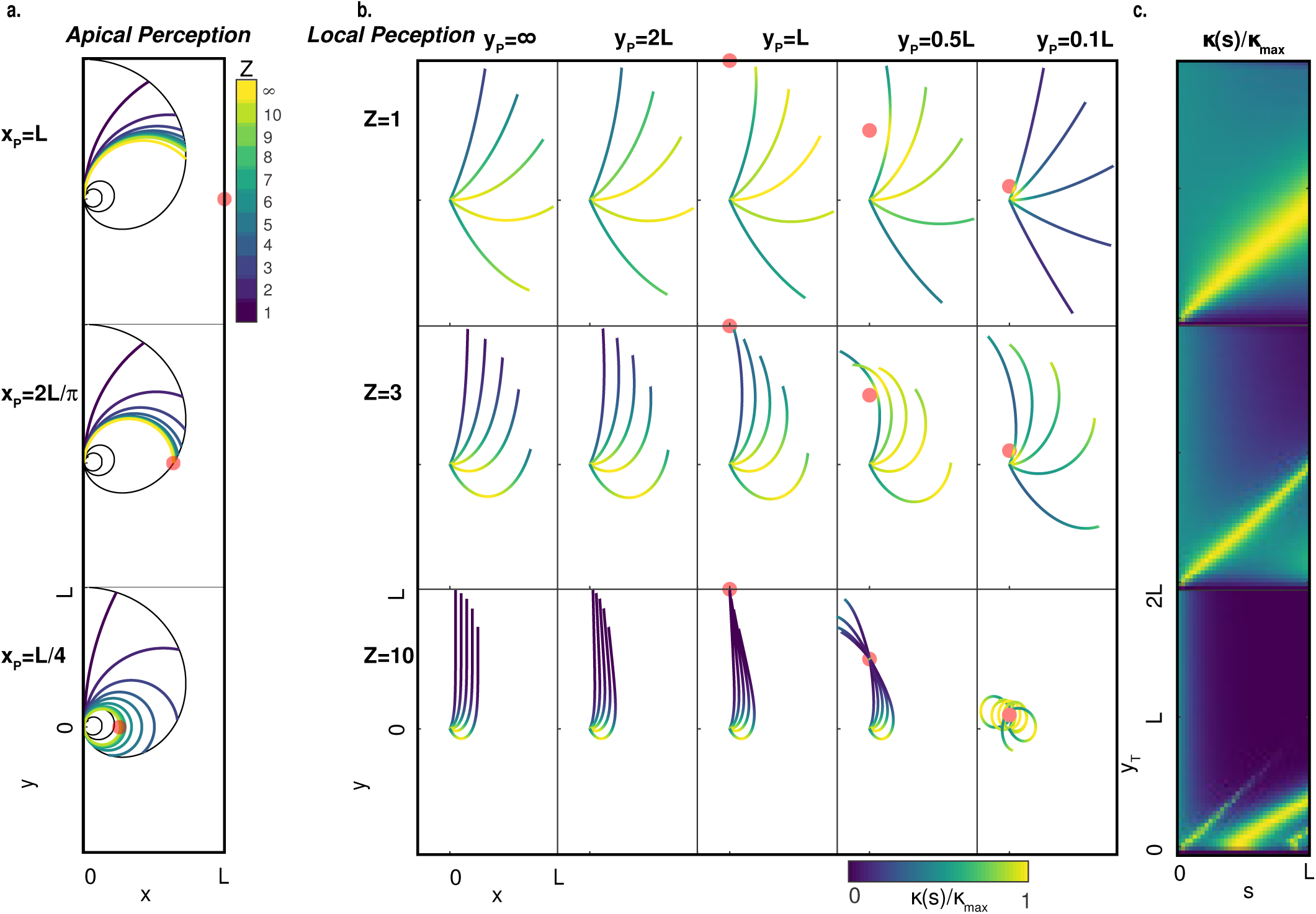
Steady state analysis of point tropism. (a) Apical sensing: Steady states of dynamics of initially straight and vertical organs, *θ*_0_ = 0, in response to a point signal (red dot) placed at different positions (*y*_*P*_ = 0 and *x*_*P*_ = *L*, 2*L/π, L/*4 from top to bottom). Colors reflect different values of *Z*. The black curves represent the possible apex positions of steady states compatible with the initial conditions, as described in the text and calculated in the SM. (b) Local sensing: Steady states for an initially straight organ clamped at different angles at the base *θ*_0_ = *kπ/*6 with *k* = 1, 2, 3, 4, 5, for *Z* = 1, 3 and 10 (from top to bottom), as a response to point signals placed at different distances from the base marked by a red dot (*x*_*T*_ = 0 and *y*_*T*_ = ∞, 2*L, L, L/*2*, L/*10 from left to right). The colors denote the local curvature normalized by the maximal possible curvature *κ*(*s*)*/κ*_*max*_ (c) Local sensing: Normalized curvature map *κ/κ*_*max*_, of the steady states for an organ clamped at the base with *θ*_0_ = *π/*6, as a function of the the position along the organ *s* and the position of the point tropism (*x*_*T*_ = 0 and 0 *< y*_*T*_ < 2*L*.

When a signal is placed outside of the trajectory of possible apex positions (top figure in Fig. 2a with *x*_*P*_ = *L* and *y*_*P*_ = 0), then we note that in the absence of postural control (proprioception), *γ* = 0 and *Z* = ∞, the organ curves and stops when the apical tip of the organ is aligned with the direction of the point signal. As *Z* decreases (proprioception increases), the organ is not able to completely align its tip with the stimulus directions, and stops before that. If the point signal intersects the line of possible apex positions (middle figure in Fig. 2a with *x*_*P*_ = 2*L/π* and *y*_*P*_ = 0), then in the absence of proprioception the apical tip will reach the point, which by definition means the tip is aligned with the direction of the signal. As before, higher proprioceptive sensitivity does not allow organs to reach the stimulus. Finally, when the point is within the trajectory of possible apex positions (bottom figure in Fig. 2a with *x*_*P*_ = *L/*4 and *y*_*P*_ = 0), the organ may loop in order to align with the point stimulus.

We now continue to the more complex case of local sensing, where sensing is no longer limited to the apex, but rather occurs along the organ as described in Eq. 1. Similarly to the apical case, we start by analyzing the steady state with signal points placed at different positions, as shown in Fig. 2b. Each row corresponds to different values of *Z*, and each column corresponds to a signal placed vertically from an organ’s base at distances *y*_*P*_ = 0.1*L*, 0.5*L, L*, 2*L* and *y*_*P*_ = corresponding to normal tropism. Each shown configuration represents the steady state for an initially straight organs *κ*(*s, t* = 0) = 0 placed at different base angles: *θ*_0_ = *kπ/*6 for *k* = 1, 2, 3, 4, 5, and the color code represents the value of local curvature along the organ normalized by the maximal curvature *κ*(*s*)*/κ*_max_, exhibiting large variations. This demonstrates that in the case of local sensing the proximity of the point signal introduces local perturbations in the curvature along the organ, associated with the variations in the perceived stimulus angle. When the point signal is infinitely distant *y*_*T*_ = *∞*, the steady state configuration corresponds to the AC model [47]. The organ curves to reach the vertical and higher values of *Z* yield a shorter convergence length *L*_*c*_. However contrary to the AC model the length of the curved zone is not constant for a given *Z*, increasing for large base angles *θ*_0_ due to the sine function.

The local effects on curvature are represented in detail in Fig. 2c, where the normalized curvature *κ*(*s*)*/κ*_max_ of the steady state configuration of an organ, initially clamped at an angle *θ*_0_ = *π/*6, is plotted as a function of position along the organ (0 ≤ *s* ≤ *L*), and position of the point signal with *x*_*P*_ = 0 and 0 ≤ *y*_*P*_ ≤ 2*L* (again for *Z* = 1, 3, 10). For large values of *y*_*P*_ the curvature is concentrated near the base and decreases exponentially towards the tip. As the point signal comes closer, with *y*_*P*_ < *L*, a peak in curvature is observed in the region closest to the signal point, in line with the observation that close to the signal point the perceived angle of the signal varies greatly (illustrated in Fig 1).

The diverse configurations resulting from local curvature perturbations mean that an analytical map of possible positions of the apical tip, as we found in the apical case, is harder to establish, and we will resort to a numerical analysis of the space of possible steady state positions of the apex for initially vertical and straight organs. We expect this to depend on both *Z* and the distance to the point signal. Let us first consider the case of an infinitely distant signal. Fig. 3a shows different trajectories of all possible apex positions for different values of *Z*, where the base of the organ is placed at the origin (0, 0). The color scheme represents the angle of the point stimulus relative to the origin *θ*_0*P*_. When *Z* = ∞ the solution is a circle of radius *L*, starting straight ahead for a signal placed in direction *θ*_0*P*_ = 0 (dark blue) and ending in the opposite direction for *θ*_0*P*_ = *π* (yellow). This is the case since with no prorioception the curvature can be infinitely concentrated near the base, and the organ aligns perfectly with any stimulus direction. As the proprioceptive sensitivity increases, and *Z* decreases, the range of possible positions is reduced, forming a closed curve. This reflects the point, that with proprioception, when a signal is placed almost directly behind the organ (high values of *θ*_0*P*_ marked in yellow), the organ is not able to bend enough and a different configuration is preferable. We note that the outer trajectory is common to all values of *Z*.

**FIG. 3.**
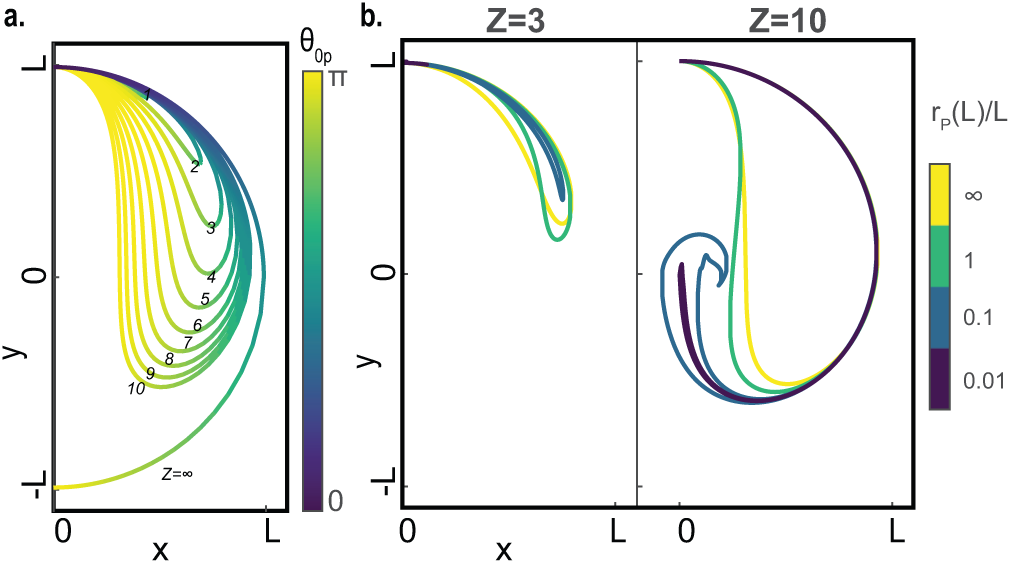
Possible steady state apex positions for local sensing. (a) Trajectories mark the steady state position of the apical tip of an initially straight and vertical organ, with local sensing, in response to a stimulus placed at an infinite distance, at various angles relative to the base of the organ *θ*_0*P*_. Each trajectory represents a different value of *Z* (marked), and the color scheme along the trajectory represents the angle at which the distant stimulus was placed relative to the origin, as detailed in the main text. (b) Simulations were run with the same initial conditions, but the point stimulus was placed in different positions relative to the organ. Each trajectory represents the steady state positions of the apical tip of all configurations where the tip stopped at a given distance *r*_*T*_(*L*)*/L* = 0.01, 0.1, 1, ∞, as also shown by color. On the left are the trajectories for *Z* = 3, and on the right *Z* = 10.

In order to study effects of the distance of the stimulus to the organ, in Fig. 3b we plot possible steady state apex positions placed at a given distance *r*_*P*_(*L*) from the stimulus, as defined in Eq. 3. Trajectories are plotted for distances *r*_*P*_(*L*)*/L* = 0.01, 0.1, 1, ∞. For *Z* = 3 the range of possible solutions does not vary significantly, while for *Z* = 10 the inner curve is affected (associated with larger angles), while only small variations are observed on the outer part, (associated wiht smaller angles). From these analyses we deduce that the outer curve for a large value of *Z* is a good approximation for the possible steady state apex positions, apart from cases of stimuli placed behind the organ at at large angles relative to the organ base. This will be instrumental in understanding the steady state configurations in the case of two interacting organs, as detailed in the next section.

## IV. ALLOTROPIC INTERACTION BETWEEN TWO ORGANS

Building on our understanding of the dynamics and steady states of organs in response to a point stimulus fixed in space, we now proceed to the case of pairs of interacting organs. We assume that the signal for allotropism is emitted at the apex. Each organ then perceives the neighbor’s signal either apically or locally. The position of the point signal is then no longer fixed in space, but rather moves with the organ’s apex. We consider two organs vertically aligned *θ*(*s*, 0) = 0, separated by a distance ∆*x* and ∆*y*, as illustrated in Fig. 1b, so that they are symmetrically positioned around the origin, *i.e.* the left organ is placed at *x*_1_(*s* = 0) = − ∆*x/*2 and *y*_1_(*s* = 0) = 0, while the right organ is placed at *x*_2_(*s* = 0) = ∆*x/*2 and is placed at some position in the y direction with *y*_2_(*s* = 0) = ∆*y*.

Following Eq. 1 we now have two equations of motion, one for each organ, coupled through the perceived angle of the stimulus emitted by each other’s tip:

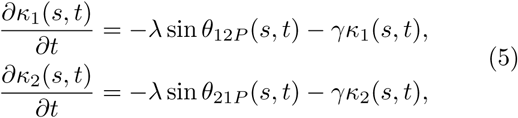

Here *θ*_1_(*s, t*) and *θ*_2_(*s, t*) are the local angles of the left and right organ respectively, and *κ*_1_(*s, t*) and *κ*_2_(*s, t*) are the respective local curvatures. We assume here that both organs have the same dynamical value *Z* (as well as *γ* and *λ*). The angle of the stimulus emitted by the tip of the right organ as perceived by the left organ is *θ*_12*P*_ (*s, t*), while *θ*_21*P*_(*s, t*) is the opposite. Following Eq. 2 they are therefore defined as

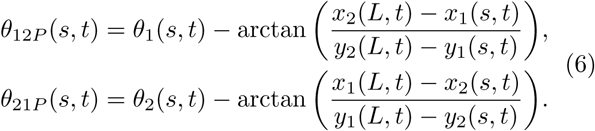

As in the previous section, we first consider apical perception, replacing the perception angles in Eq. 6 with *θ*_12*P*_(*L, t*) and *θ*_21*P*_(*L, t*). We simulated pairs of organs placed at different relative positions, 0 ≤ ∆*x* ≤ 2.3*L* and −2.3*L* ≤ ∆*y* ≤ 2.3*L*, where negative values of ∆*x* just yield mirror configurations. For each simulation we take note of the apex position of the left organ in its steady state configuration, *x*_1_(*s* = *L*) and *y*_1_(*s* = *L*), and Fig. 4a plots these values for *Z* =∞, as a function of relative organ position ∆*x* and ∆*y*. The values of the right organ *x*_2_(*L*) and *y*_2_(*L*) for a given value of ∆*y* are equivalent to the values of *x*_1_(*L*) and *y*_1_(*L*) for *−*∆*y*, since these are identical flipped configurations. These maps can be considered as the *phase space* of steady state configurations of pairwise interactions, and will be useful in our understanding, particularly in the more complex case of local sensing.

**FIG. 4.**
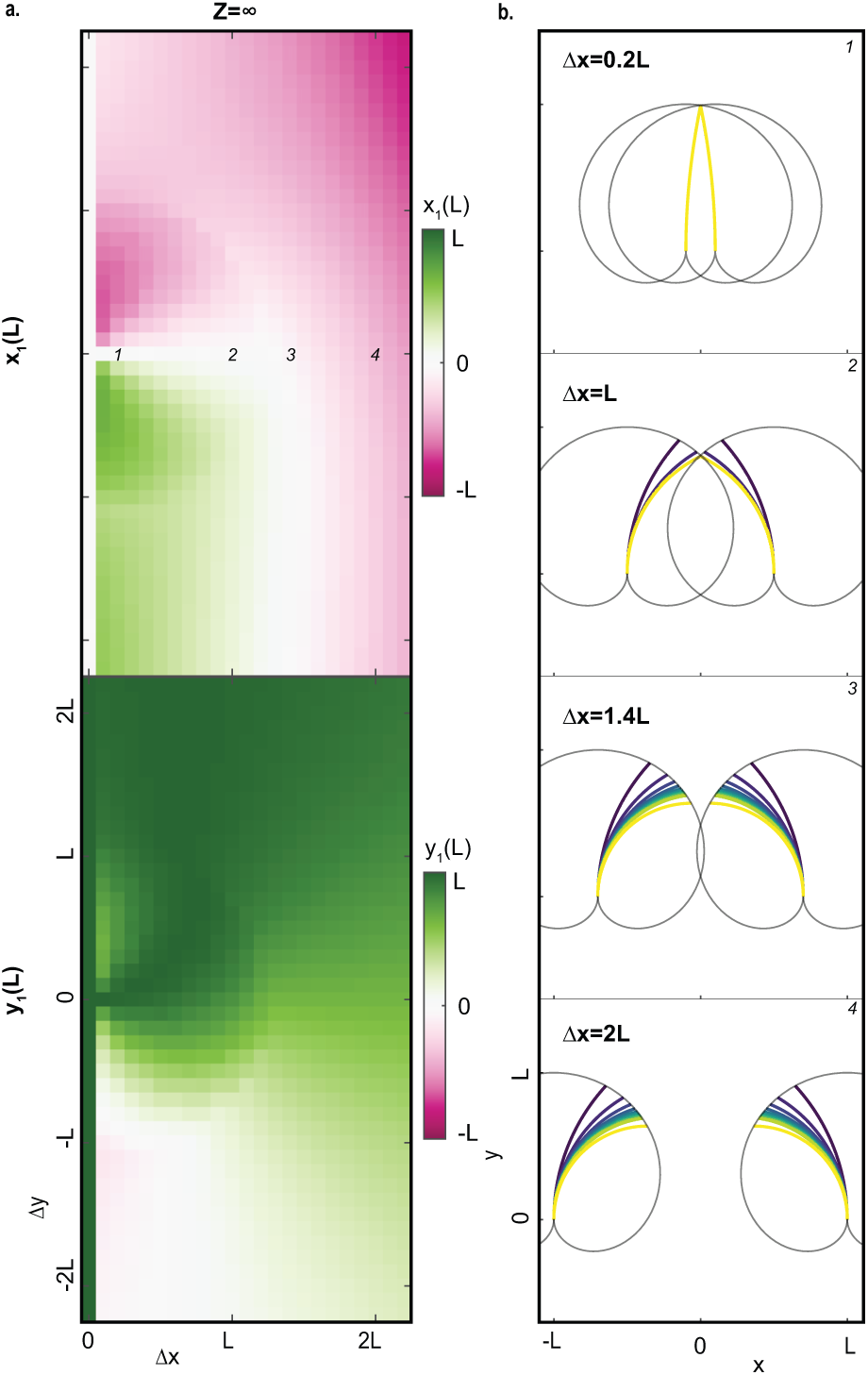
Steady states of allotropic interaction with apical perception. Simulations of pairs of initially straight organs *κ*_1_(*s*, 0) = *κ*_2_(*s*, 0) = 0 placed at different relative distances according to the geometry described in Fig. 1b. Each organ apically senses the stimulus emitted from the apex of its neighbor, with no proprioception, *Z* = ∞. Values of the steady state tip positions *x*_1_(*L*) and *y*_1_(*L*) for different relative base-to-base positions (∆*x*, ∆*y*) are represented by color in the top and bottom plots respectively. (b) Steady states of four different symmetric configurations, with organs placed at distances ∆*x* = 0.2*L, L*, 1.4*L*, 2*L* from top to bottom, marked respectively in the phase space in (a). Colors represent difdifferent values of *Z*, with the same colorbar as in Fig. 2a. The black trajectories mark the possible steady state apical positions calculated in the SM.

Fig. 4b displays configurations of simulations from four different points in the phase space in the left panel, namely ∆*x* = 0.2*L, L*, 1.4*L*, 2*L*, with ∆*y* = 0 for all (i.e parallel organs). The different colors correspond to different values of *Z*. Following Fig. 2a the organs are overlaid on the trajectory of possible apex positions in steady state. When the organs are far away (∆*x* = 2*L*), and in the absence of proprioception *Z* = ∞, the curvature of both organs is such that their apices are aligned to each other, *i.e. θ*_12*P*_ = *θ*_21*P*_ = 0, the perceived stimulus is zero, and the organs stop moving. Similarly to Fig. 2a, organs with proprioception stop to curve before their apical part are aligned. This is reflected in the associated point in phase space, with *x*_1_(*L*) *<* 0 (the apex of the left organ does not pass the center line between the organs), while *y*_1_(*L*) *>* 0 (the apex is in the upper quadrant). Since the organs are placed in parallel, they are the mirror image of each other, and therefore *x*_2_(*L*) *>* 0 and *y*_1_(*L*) *>* 0.

From Fig. 4b we see that at some relative distance ∆*x* the curves of possible apex positions will intersect. We can calculate this distance by considering that the first point of contact will be at the outermost point along the curve, which is reached by an apex in the case that the two apices are aligned with each other, with *Z* = ∞. In this case the apices are horizontal, and therefore the organs cover a quarter of a circle. In this case the distance along the x-axis from the organ base to the apex position *x*_1_(*L*) − *x*_1_(0) is just the radius of curvature of the organ, given by *ρ* = 2*L/π*. Hence when organs are placed at smaller distances, the trajectories of their possible steady state apical positions intersect. For ∆*x >* 4*L/π* they align towards each other before reaching the intersection point, and stop there (as shown for ∆*x* = 1.4*L*). For ∆*x <* 4*L/π* they reach this intersection point before they are able to align, where they touch. If a point signal is placed at the apex, then by definition the apex is aligned towards it, and again the perceived signal is zero. Hence the organs will stop at the intersection point, as shown for ∆*x* = *L* and ∆*x* = 0.2*L*. Organs with apical sensing do not overshoot the desired solution in the direction of the stimulus [48], and it is therefore expected that the organs will indeed stop at the first point where the signal is minimized.

The case of local perception provides a more complex picture due to the local curvature perturbations discussed earlier for single organs in proximity of a point stimulus. Following the analysis for apical sensing, we plot in Fig. 5 the phase space represented by *x*_1_(*L*) and *y*_1_(*L*) of steady state configurations, for both *Z* = 3 and *Z* = 10. For *Z* = 3 the phase space is smooth, and solutions are continuous. Comparing the phase space with that for apical sensing, we note that the picture is qualitatively similar, producing very similar dynamics. For *Z* = 10 the picture is dramatically different. First, we observe that at certain relative positions no convergence is observed, represented in black. In these cases both organs oscillate without reaching a steady state, which will be discussed later. Furthermore, the phase space is no longer continuous, exhibiting discontinuous jumps between phases, reminiscent of a first-order phase transition. The phases are distributed with a roughly circular geometry.

**FIG. 5.**
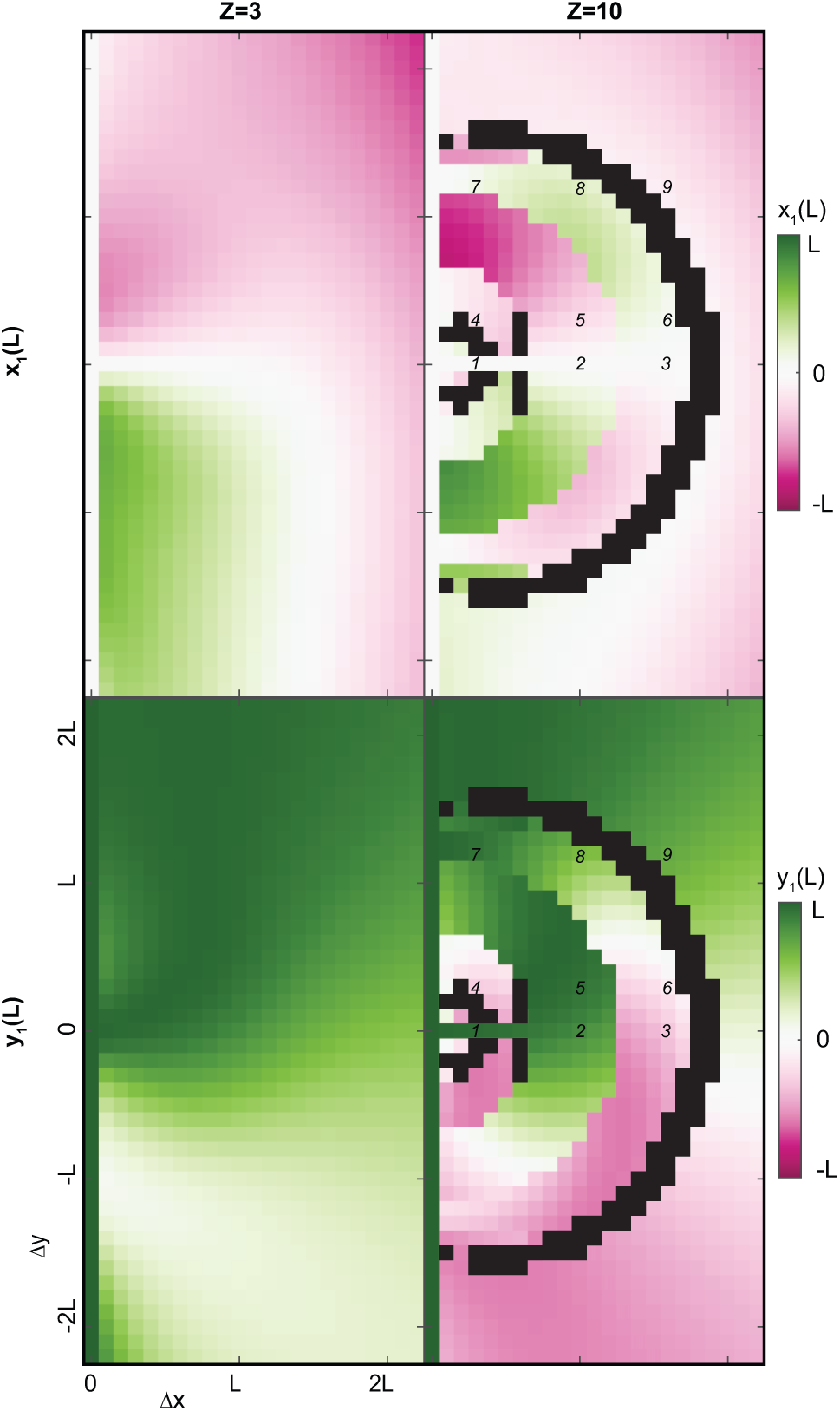
Steady state phase space of allotropic interaction with local perception. Simulations of pairs of initially straight organs placed at different relative positions. Positions of the apical tip of the left organ at steady state, *x*_1_(*L*) (top panel) and *y*_1_*L*) (bottom panel), as a function of ∆*x* and ∆*y*. The left side describes the phase space for *Z* = 3 and the right *Z* = 10. Black represents positions where dynamics do not converge to a steady state. Examples of specific configurations from different places in the phase space, as marked by numbers, are shown in Fig. 5

Following the analysis for apical sensing, we present representative steady state configurations from different areas in the phase space in Fig. 6. We note that the organ configurations are overlaid on the common outer trajectory of possible steady state positions of apices, as discussed in Fig. 3, which is a good approximation as long as signals are not placed behind the organ base. While this approximate trajectory does not represent the full picture, it does provide some intuition when discussing the different types of steady states, or phases.

**FIG. 6.**
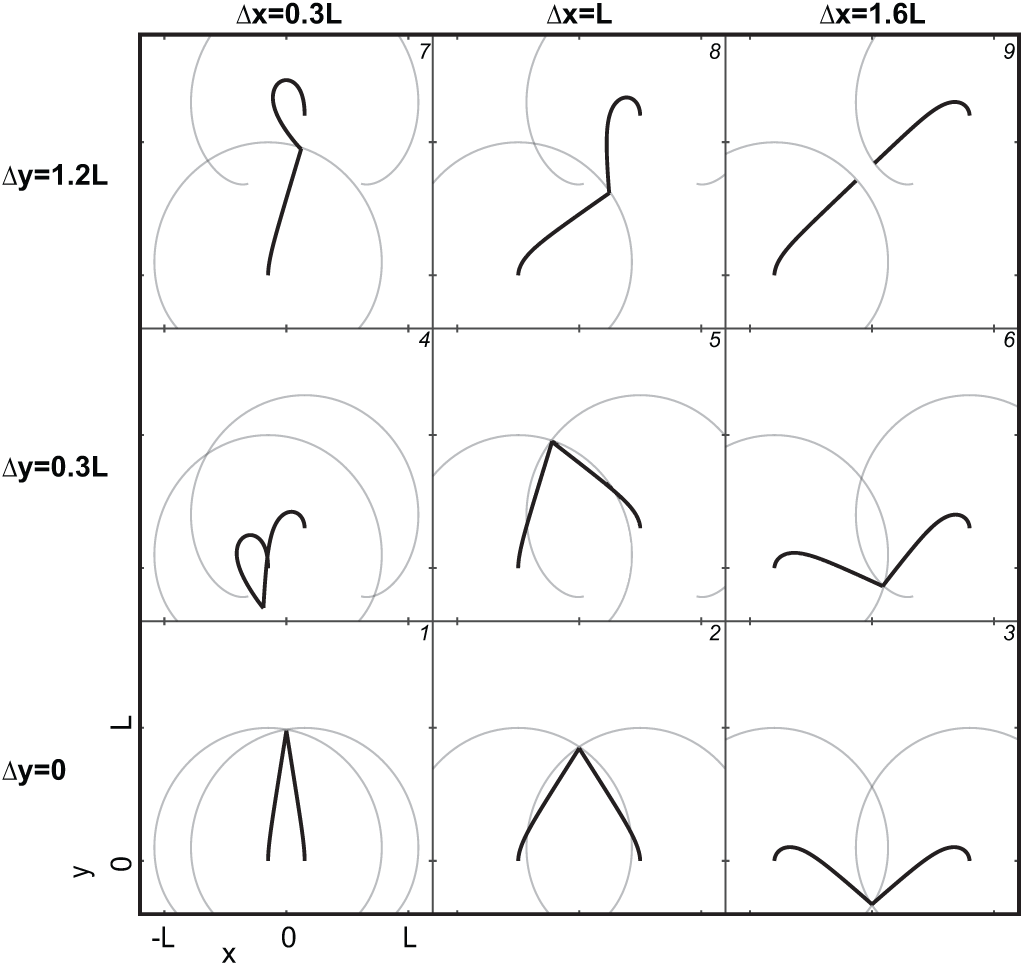
Examples of steady state configurations of pairs of interacting organs with local perception. Organs are placed at different base-to-base distances sampling the phase space in Fig. 5. Simulations were run with *Z* = 10, with ∆*x* = 0.3*L, L*, 1.6*L* from left to right, and ∆*y* = 0, 0.3*L*, 1.2*L* from bottom to top. The organs are overlaid on the approximated curve of possible steady state apex positions (which is common to all distances, as shown in Fig. 3b). Each configuration is numbered and marked on the respective position in the phase space in Fig. 5.

Going back to the phase space in Fig. 5, the outermost area, defined for organs whose bases are placed at a distance roughly larger than 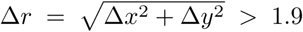, represents the simplest case of organs placed at large distances. The organs are too far away to intersect or align, and this is equivalent to the pattern observed when *Z* = 3. An example of a relevant steady state configuration is shown in plot (9) of Fig. 6.

Next we consider the phase defined for distances roughly within 0.6*L <* ∆*r <* 1.2*L*. At these distances the apex trajectories intersect, as can be seen in examples (2) and (5) in Fig. 6, and the apices touch at the first intersection point. Let us consider the symmetric cases where ∆*y* = 0 (*e.g.* configuration(2)). We see in the phase space that *x*_1_(*L*) = *x*_2_(*L*) = 0, *i.e.* the tips touch at the center-line, while *y*_1_(*L*) *>* 0 and *y*_2_(*L*) *>* 0. Since the values of *x*_1_(*L*) and *y*_1_(*L*) change in a continuous manner, we can assume that in all configurations within this range the organ tips converge at the first intersection point.

We now proceed to consider the phase defined for organs placed at distances approximately within 1.2*L <* ∆*r <* 1.7*L*. Also within this range the apex trajectories intersect, as can be seen in examples (3) and (6) in Fig. 6. We note that for examples (7) and (8), which belong to this phase, the curves are no longer a good approximation due to the large angles, and a more detailed approximation is required. Looking at the phase space we see that for values of about |∆*y| <* 0.5, we have *y*_1_(*L*) *<* 0 and *y*_2_(*L*) *<* 0, *i.e.* both tips point down, clearly shown in examples (3) and (6). We note that the apices touch at the *bottom* intersection point a phase which was not observed in previous cases. This means that during their dynamics the organs did not stop at the first intersection point they encountered, but continued to the next. This is true also true of the other examples in this phase, (7) and (8). As argued before, since the values of *x*_1_(*L*) and *y*_1_(*L*) change in a smooth manner within this range, we can assume that in all configurations within this range the organ tips converge at the second intersection point.

Finally for very short distances, roughly ∆*r <* 0.5*L* we see a more complex behavior. For symmetric cases the tips touch at the first intersection point as in example (1). Upon breaking the symmetry they no longer stop at the first intersection point, and continue to the second, as shown in example (4).

In order to understand why in certain configurations organs do not stop at the first intersection point, but rather continue to the second, we need to discuss the dynamics of the organs. The fact that sharp *first-order* phase transitions are observed at roughly fixed distances, suggests that the base-to-base distance is a dominant factor. Fig. 7a shows snapshots of the dynamics of four symmetric configurations from different points in phase space, where the evolution of time is represented by color code. Fig. 7b shows the locally perceived angle along the organs, and Fig. 7c shows the local change in curvature. Let us consider the first example of two organs placed at ∆*x* = 1.2*L* apart. we observe that after leaving their initially straight conformation the organs cross each other, and exhibit damped oscillations as they converge to the final steady state configuration. This is related to the overshooting behavior found to be characteristic of local sensing organs with low proprioception (high values of *Z*) in regular tropic models [47]. Indeed with no proprioception (*Z* = ∞), it was shown that when a stimulus is placed in a perpendicular direction from the organ, as the organ approaches the direction of the stimulus the basal part continues to curve. The organ overshoots the stimulus direction, then crosses back, overshooting it in the opposite direction etc., yielding oscillations which go on indefinitely, while their amplitude decreases with time. Considering the dynamics of the organs placed at ∆*x* = 1.2*L*, we observe that when the organs overshoot they cross each other upwards, so that the apices are above the rest of the organs. The apices therefore *pull* the organs back up towards the first intersection point.

**FIG. 7.**
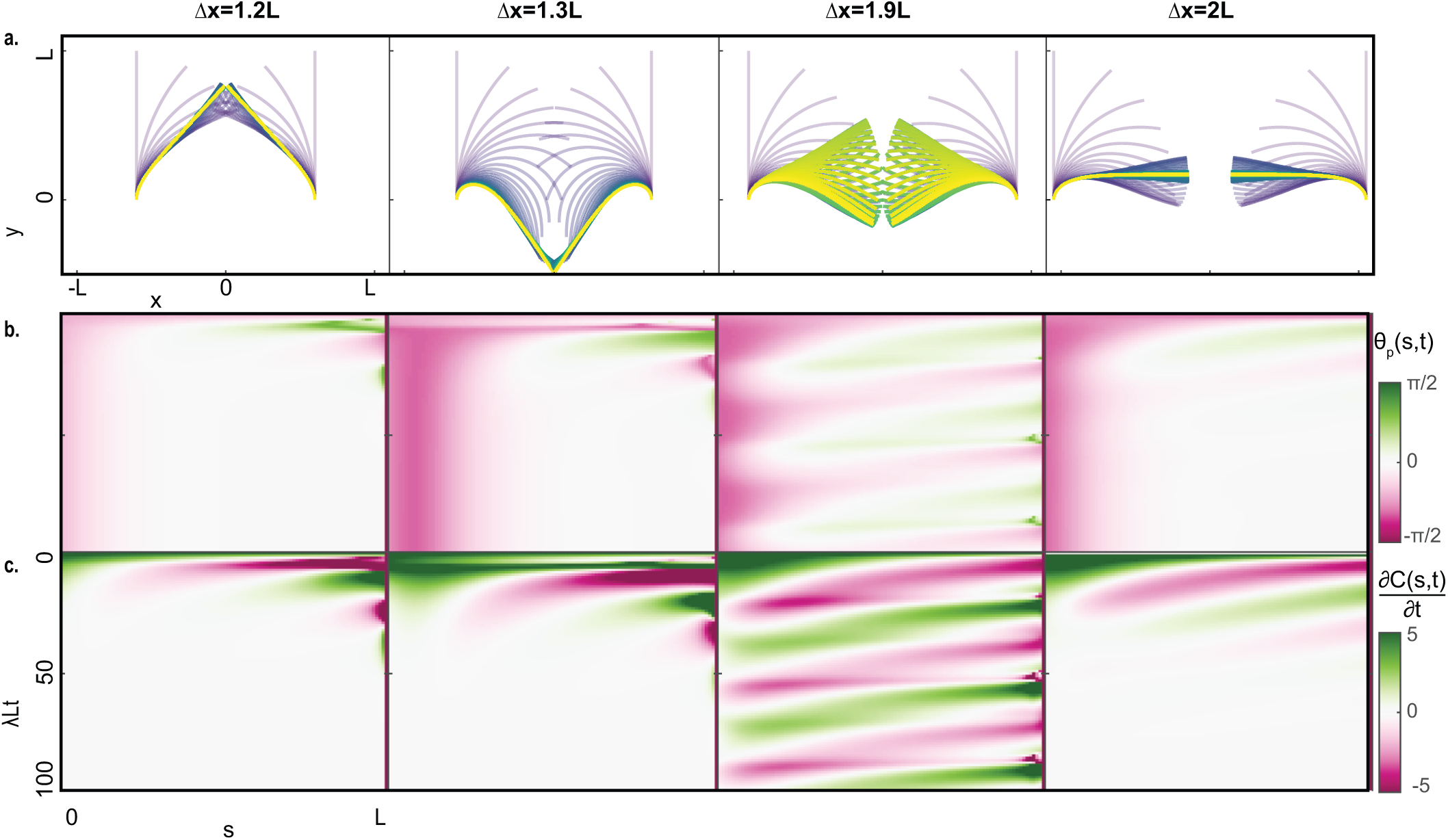
Dynamics of allotropic interactions with local perception. (a) Snapshots of dynamics of symmetric pairs of organs placed at distances ∆*x* = 1.2*L*, 1.3*L*, 1.9*L*, 2*L* from left to right. These cases are also marked in the phase space in Fig. 5. (b) Map of perceived angle *θ*_*P*_(*s, t*) as a function of *s* along the organ, and normalized time *λLt*. (c) Map of local curvature variation 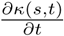 along the the organ, as a function of normalized time *λLt*.

However for larger distances, such as ∆*x* = 1.3*L*, by the time they cross each other, which is after the first intersection point, they have already curved enough so that they cross each other downwards, with the apices below most of the organ, thus *pulling* the organs towards the next intersection point. This is illustrated in Fig. 7b and c, where for ∆*x* = 1.2*L* the perceived angle goes from negative to positive values after a single time step, translated to changes in the direction of curvature propagating from the tip. For ∆*x* = 1.3*L* the change in perceived angle occurs much later. We note that this argument suggests that the form of the phase space also depends on intial conditions, or the history of the system dynamics. In the SM we show an example of organs placed at ∆*x* = 1.68*L*, but with an initial configuration placed at the first intersection point. Instead of continuing to the next intersection point as expected from the phase space, they remain stable at the first intersection, since the organs did not cross each other downwards. It is reasonable to extrapolate this for all configurations in the respective phases, however proving this is beyond the scope of this paper.

Lastly Fig. 7 also shows an example of the non-converging oscillating dynamics, associated with the black colored outer ring in Fig. 5, defined roughly for 1.7*L <* ∆*r <* 1.9*L*. Within this range the apices are extremely close to each other, but can barely touch. Fig. 7b and c illustrates that in this particular configuration, as opposed to the stable dynamics, the value of the perceived angle propagates from the apex all the way to the base. Namely, at every point in time the perceived angle at the base and at the tip are in opposite directions, directly translated to curvature changes in opposite directions (this can be seen by drawing a straight line at any time in the respective *θ*_*P*_(*s, t*) and 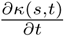 map). We note that this is a result of the proximity of the signal point to the organ, which leads to large variations in the locally perceived angle and local curvature. This is further exemplified in the SM where we show how oscillations can be initiated as a result of this geometrical argument.

## V. DISCUSSION AND CONCLUSION

In this paper we addressed the concept of collective or swarming dynamics in systems where movement is not due to self-propulsion, but rather due to growth. The goal of the paper is to develop a mathematical framework which will allow to rigorously investigate this phenomenon. Here we take inspiration from plant tropisms the growth-driven response of plants to external stimuli such as light and gravity, where a robust mathematical model has recently been developed. However the resulting framework is applicable to other systems which move by growing, including biological systems such as neurons and fungi, and robotic systems [54–56].

Within this work we have introduced two new concepts related to plant tropisms: point tropism and *allotropism*. Point tropism refers to a nearby signal source whose signal propagates radially and therefore affects different point on an object from different angles. This is in contrast to regular tropism, where the signal source is generally assumed to be far away (gravity, the sun), yielding a parallel signal field which affects objects from the same angle. This scenario is relevant for tropisms towards localized sources, for example plant roots searching for water or nutrients. We analyzed the dynamics of a single organ responding to a fixed point both for the case of apical sensing and local sensing. We were able to analytically characterize the possible configurations in the simpler case of apical sensing. However we found that in the case of local sensing, where sensing occurs along the whole organ, the proximity of the signal leads to local perturbations in the curvature of the organ, yielding complex dynamics and configurations. We made a numerical analysis which allowed to make an approximation of possible steady state configurations.

We termed *allotropism* the observed growth-driven responses of plant organs in the direction of their neighbors, for example a root growing towards or away from a neighboring root. Here we assume that a pair of organs can sense a signal emitted from a point fixed at their neighboring organ’s tip, thus coupling the point tropism dynamics of both organs. We ran simulations for pairs of organs placed at various relative positions in space, both in the case of apical and local sensing, and considered the phase space as represented by the position of the apices in the steady state configuration. We found that in the case of apical sensing the phase space is continuous, and the different types of configurations can be easily understood from the point-tropism analysis for single organs. However for local sensing we found that proprioception, the resistance of the organ to bending, has a significant effect. For high proprioceptive values the phase space is qualitatively very similar to that of the apical sensing. However for low proprioceptive values organs overshoot the direction of the stimulus, and this yields new steady state conformations which were not possible otherwise, including non converging dynamics. Indeed the phase space now exhibits discontinuities between phases roughly correlated to the base-to-base distance between the organs. We found that the possible steady state positions are determined by the intersection points of the approximated curves of possible steady state apex positions of each organ (a property of the single organ). We suggest that the sharp transitions between these phases are due to the tendency of organs to overshoot their stimulus at high values of *Z*, which causes them cross each other, effectively pulling each other towards one of a few possible steady state positions. This argument also suggests that the form of the phase space should depend on the initial conditions, which we verified in the SM.

Lastly we also observed non converging dynamics associated with self-sustained oscillations of the organs. These oscillations occur when the organs are placed at a distance such that the value of the perceived angle propagates from the apex all the way to the base, and the perceived angle at the base and at the tip are in opposite directions. This is a result of the proximity of the signal point to the organ.

The analysis brought here focused on the case of constant attractive interactions, however the interactions can easily be replaced with a repulsive interaction, or a more complex distance-dependent interaction, such as those generally used in common models for collective behavior [18, 24, 57]. The case of constant repulsion is trivial since organs will always align in opposite directions, and the signal will never be close enough to cause local curvature perturbations.

Future work may include many possible generalizations. Here we focused on the limiting case where the elongation rate of the organ is negligible compared to the bending dynamics. However this can be generalized to explicitly account for growth following previous work 2014. In addition, elasticity [58] and more complex integration of stimuli [51] can be integrated, as well as a generalization to 3D [50]. Moreover, this model assumes the signal is emitted from a point at the tip, however it is possible to consider the signals emitted from a more extended part of the organ, by assuming the signal is additive.

Lastly, this work brought an analysis of the phase space of possible dynamics and steady states of pairs of interacting organs. The mathematical framework presented here allows to take the next step and investigate collective behavior or swarming of multiple organs. The ideas presented here will hopefully spark interest in the concept of collective behavior in a class of systems with inherently different dynamics, and are relevant both for researchers working on biological systems where movement is due to growth, as well as for the growing community of plant-inspired robotics.

## ACKNOWLEDGMENTS

This work was performed in part at the Aspen Center for Physics, which is supported by National Science Foundation grant PHY-1607611. Research was partially supported by the Israel Science Foundation grant (1981/14).

## VI. SUPPLEMENTARY MATERIAL

### A. Steady state form of point source tropism with apical sensing

In what follows we calculate analytically the form of the steady state. If the stimulus is sensed only at the apex, the dynamics follow:

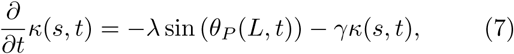

where 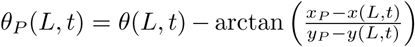 is the relative angle between the apex of the organ at point *s* = *L* at time *t* and the point signal at (*x*_*P*_,*y*_*P*_).

When sensing is apical, the whole growth zone responds identically to the same signal leading to a constant curvature *κ* along the organ, associated with a radius of curvature by the geometrical relation *R* = 1*/κ*. The apex therefore moves along a track corresponding to the endpoints of all possible arcs of a constant length *L*, as shown in Fig. 8.

**FIG. 8.**
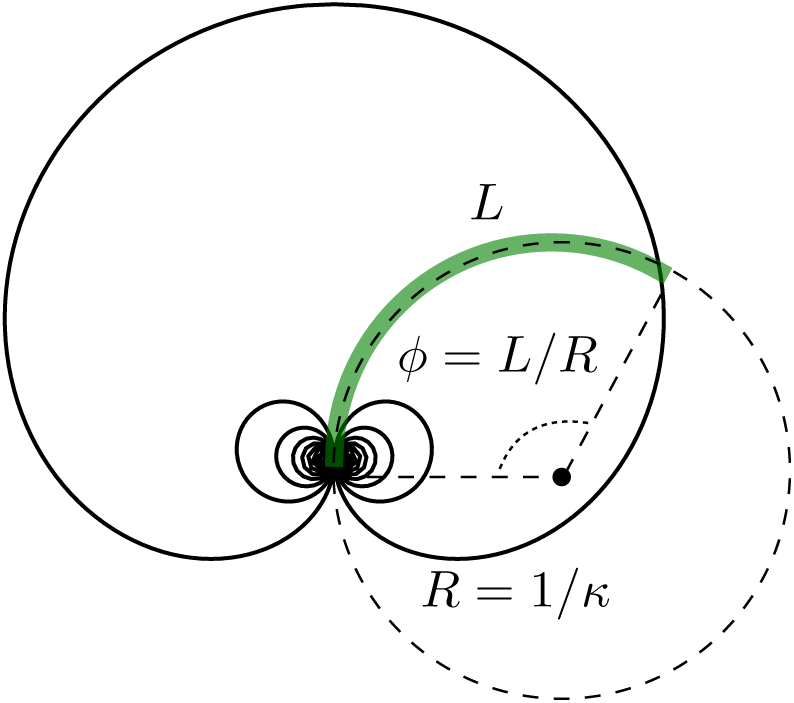
Possible apex locations for apical sensing. The red line represents the organ of length *L* and constant curvature *κ*, whose base is clamped at the bottom. The dotted line represents the imaginary circle completing the arc of the organ, with a radius of curvature *R* = 1*/κ*, and the tangential angle *φ* which spans the arc is *φ* = *L/R* = *κL*.

For an organ with a given curvature *κ* we consider the imaginary circle completing the arc of the root, illustrated as a dashed line in Fig. 8. The radius of curvature is *R* = 1*/κ*, and the tangential angle spanning the arc is therefore *ϕ* = *L/R* = *κL*. The position of the apex for an organ with a given fixed curvature *κ* can then be written as:

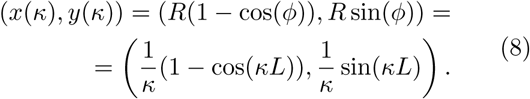

This parametrization provides the track of all possible positions of the organ, plotted as a black solid line in Fig. 8.

The curvature of the steady state form of the organ can be found by substituting this parameterization and 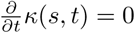 in Eq. 7, yielding:

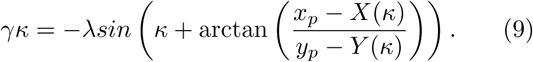

This can be generalized to multiple point signals, following Eq. **??**. It is useful to note that in the case of no proprioception *γ* = 0, the steady state solution yields *θ*_*P*_(*L*) = *nπ*, meaning that the apex of the organ is aligned with the direction of the point source.

### B. Oscillations result from signal proximity, and uctuations in locally perceived signal angle

The oscillations are the consequence of an apparent propagation of the value of perceived angle from the apex to the base that is correlated with the curvature variation. We simplify the geometry of the system in order to reveal the minimal requirements for oscillations. We consider the case of two organs facing each other so that *θ*_01_ = *π/*2 while *θ*_02_ = − *π/*2 and separated with a distance ∆*x* = (2 + *ϵ*_*x*_)*L*, so the organs are close but not intersecting. Fig. 9 shows this setup with *ϵ*_*x*_ = 0.01. In this perfect mirror symmetry, the system is at the steady state, and no movement is observed. We then make a small perturbation, and tilt both organs at their base with a small angle deviations ∆*θ* = ± *ϵ*_*θ*_, so that *θ*_01_ = *π/*2 − *ϵ*_*θ*_ and *θ*_02_ = − *π/*2 + *ϵ*_*θ*_, in order to keep the mirror symmetry. This very constrained set of symmetries and initial conditions is sufficient to generate oscillations, shown in Fig. 9, which are amplified by removing proprioception, taking *Z* = ∞ for *γ* = 0. The plot of the locally perceived stimulus angles illustrates that at any point in time the base and tip sense opposite directions, translated to opposite directions of motion. Since these oscillations are not damped, the system finally reaches self-entertained oscillations.

**FIG. 9.**
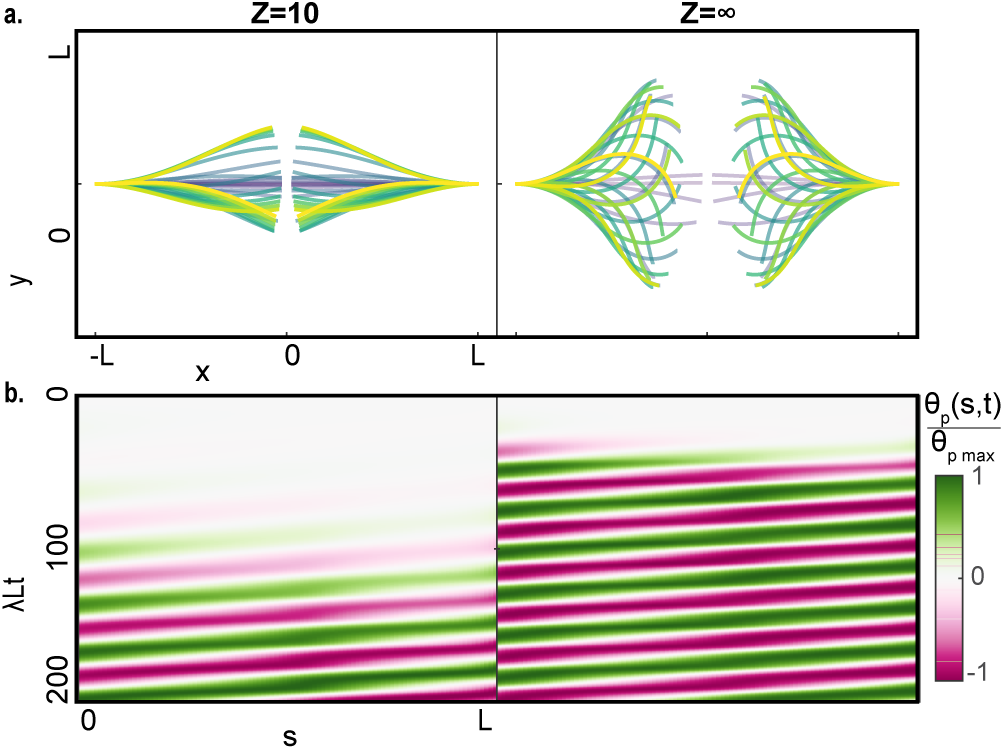
Oscillatory dynamics of allotripic interactions with local perception. (a) Snapshots of oscillatoriy dynamics in the case of two organs which are initially straight and facing each other in a mirror configuration *θ*_1_(0) = *π/*2 − *ϵ*_*x*_ and *θ*_2_(0) = −*π/*2 + *ϵ*_*x*_. *ϵ*_*x*_ = 0.01 is a small perturbation to break the symmetry of the system. The two bases are separated by a distance ∆_*y*_ = 2.01*L* so that the two organs cannot intersect. The dynamics are computed for *Z* = 10, on the left, and in absence of proprioception, *Z* = *∞* on the right. (b) Map of the perceived angle along the organ, normalized by the maximal perceived angle observed in the simulation, *θ*_*P*_(*s, t*)*/θ*_*P*_ max, as a function of the normalized time *λLt*.

### C. Stable state depends on intial conditions

Fig. 10 shows that starting with initial conditions at the first intersection point will result in a steady state at that point rather than the second intersection point as expected for initially vertical organs. Here we start from the simulation of initially straight and vertical organs placed ∆*x* = 1.2*L* from each other (shown in Fig. 7). We then start a new simulations where the distance between the organs is increased by a small increment of 0.01*L*, and the initial configuration is the steady state configuration of the previous simulation (touching at the first intersection point of the apex position curve). We continue this process and find that this steady state configuration, with both organs touching at the first intersection point, remains stable well into the distance range where the stable state was at the second intersection point (for simulations starting from initially vertical organs as in the main text). This is shown in the left panel of Fig. 10 for ∆*x* = 1.68*L*. However when this process is continued to ∆*x* = 1.7, the configuration is such that the organs start to oscillate and overshoot, and therefore cross each other downwards, and move to a new steady state at the second intersection point of the possible apical positions. We note that this distance is indeed borderline with the non converging phase. This observation is in line with the argument that the precursor to the transition from the first intersection point to the second is an overshoot of the organs where they cross each other downwards, pulling each other to the second intersection. This also shows the dependence of the phase space on the initial conditions.

**FIG. 10.**
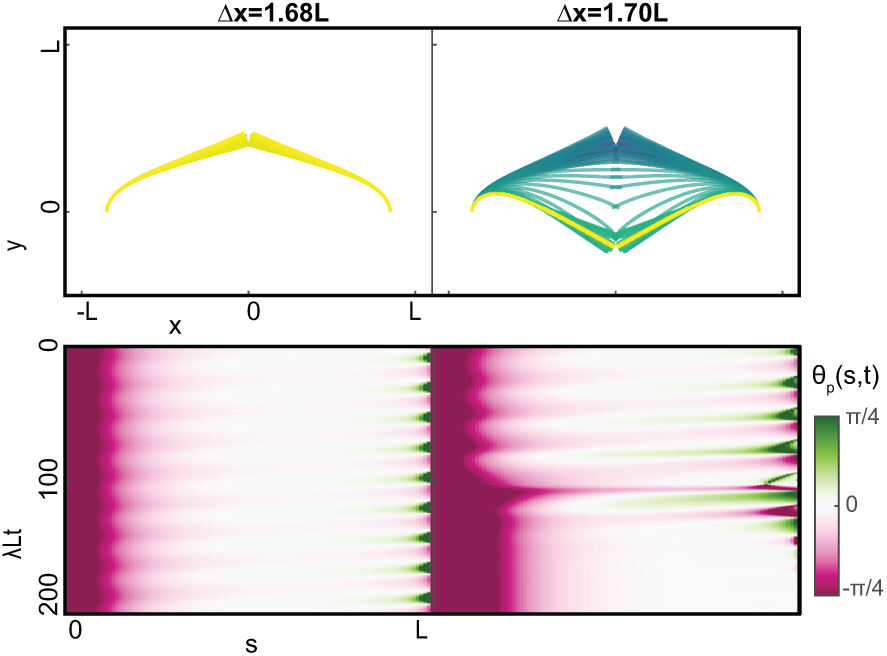
Identification of transition between steady states for allotropic interactions with local perception. Here we start from the simulation of initially straight and vertical organs placed ∆*x* = 1.2*L* from each other, and use the steady state as the initial condition for a new simulation, where the distance between the organs is increased by a small increment of 0.01*L*. Following this procedure, the left panel shows the dynamics at ∆*x* = 1.68, while the right panel shows the dynamics at ∆*x* = 1.7.

